## Supplementary material for "Group identification drives brain integration for collective performance": SM

**Table S1.** MNI coordinate Position of 3×5 optode probe set

|  | **MNI coordinate Position** | | |  |  |
| --- | --- | --- | --- | --- | --- |
|  | **x** | **y** | **z** | BrodmanArea (Chris rorden' MRIcro) | **Percentage** |
| **CH01** | 36 | 40 | 42 | 9 - Dorsolateral prefrontal cortex | 0.967742 |
| **CH02** | 13 | 50 | 46 | 9 - Dorsolateral prefrontal cortex | 1 |
| **CH03** | -11 | 50 | 45 | 9 - Dorsolateral prefrontal cortex | 1 |
| **CH04** | -35 | 40 | 42 | 9 - Dorsolateral prefrontal cortex | 0.913043 |
| **CH05** | 47 | 42 | 28 | 45 - pars triangularis Broca's area | 0.94382 |
| **CH06** | 26 | 57 | 33 | 46 - Dorsolateral prefrontal cortex | 0.504854 |
| **CH07** | 2 | 59 | 34 | 10 - Frontopolar area | 0.934211 |
| **CH08** | -23 | 56 | 33 | 6 - Dorsolateral prefrontal cortex | 0.654762 |
| **CH09** | -45 | 42 | 27 | 45 - pars triangularis Broca's area | 0.92233 |
| **CH10** | 38 | 19 | 58 | 46 - Dorsolateral prefrontal cortex | 0.789474 |
| **CH11** | 14 | 68 | 23 | 10 - Frontopolar area | 1 |
| **CH12** | -13 | 67 | 22 | 10 - Frontopolar area | 1 |
| **CH13** | -35 | 58 | 19 | 46 - Dorsolateral prefrontal cortex | 0.916667 |
| **CH14** | 47 | 53 | 2 | 46 - Dorsolateral prefrontal cortex | 1 |
| **CH15** | 27 | 68 | 8 | 10 - Frontopolar area | 0.934211 |
| **CH16** | 2 | 68 | 9 | 10 - Frontopolar area | 1 |
| **CH17** | -24 | 68 | 8 | 10 - Frontopolar area | 0.972973 |
| **CH18** | -45 | 53 | 1 | 46 - Dorsolateral prefrontal cortex | 1 |
| **CH19** | 38 | 63 | -7 | 10 - Frontopolar area | 0.622222 |
| **CH20** | 15 | 71 | -3 | 11 - Orbitofrontal area | 0.615385 |
| **CH21** | -13 | 71 | -3 | 11 - Orbitofrontal area | 0.770833 |
| **CH22** | -36 | 63 | -7 | 10 - Frontopolar area | 0.586207 |

**Table S2.** MNI coordinate Position of 2×4 optode probe set

|  | **MNI coordinate Position** | | |  |  |
| --- | --- | --- | --- | --- | --- |
|  | **x** | **y** | **z** | **BrodmanArea (Chris rorden' MRIcro)** | **Percentage** |
| **CH01** | -34 | 2 | 66 | 2 - Primary Somatosensory Cortex | 0.65079 |
| **CH02** | -45 | -26 | 67 | 22 - Superior Temporal Gyrus | 0.48867 |
| **CH03** | -46 | -54 | 59 | 39 - Angular gyrus, part of Wernicke's area | 0.67845 |
| **CH04** | -41 | 14 | 59 | 43 - Subcentral area | 0.83019 |
| **CH05** | -51 | -14 | 58 | 22 - Superior Temporal Gyrus | 0.75851 |
| **CH06** | -57 | -43 | 54 | 37 - Fusiform gyrus | 0.44554 |
| **CH07** | -57 | 43 | 54 | 19 - V3 | 0.8 |
| **CH08** | -54 | 2 | 50 | 21 - Middle Temporal gyrus | 0.54808 |
| **CH09** | -64 | -27 | 45 | 21 - Middle Temporal gyrus | 0.47187 |
| **CH10** | -63 | -51 | 37 | 37 - Fusiform gyrus | 1 |

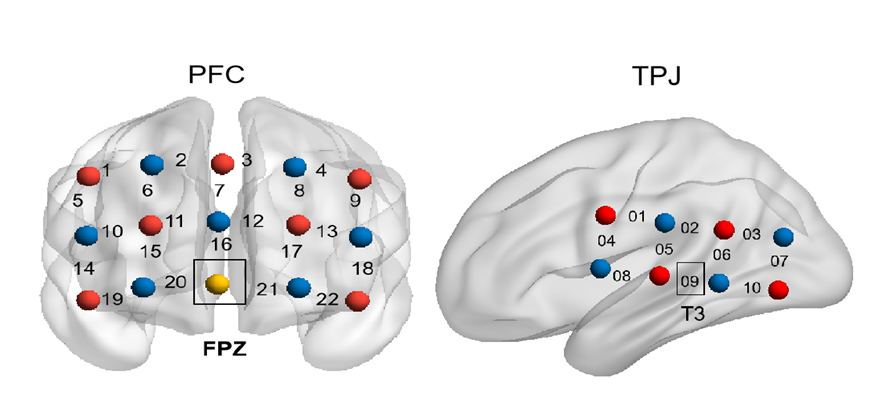

**Figure S1.** Probe location and measure the brain activity simultaneously. Optode probe sets. The sets were placed over the prefrontal and left TPJ regions.

**The results of HbR.**

We attempted to verify if the pattern of associated results was comparable to that of HbO when the analyses of HbR were conducted. First, by performing one-sample *t*-tests for GNS, we observed a significantly increased GNS in the OFC (CH20, *t* = 2.11, *p* = 0.030, FDR corrected; CH21, *t* = 6.76, *p* < 0.001, FDR corrected). We then conducted independent *t*-test on GNS in OFC (CH21), indicating a significant difference between the High and Low Group Identification groups (*t*_58_ = 2.04, *p* = 0.040, FDR corrected).

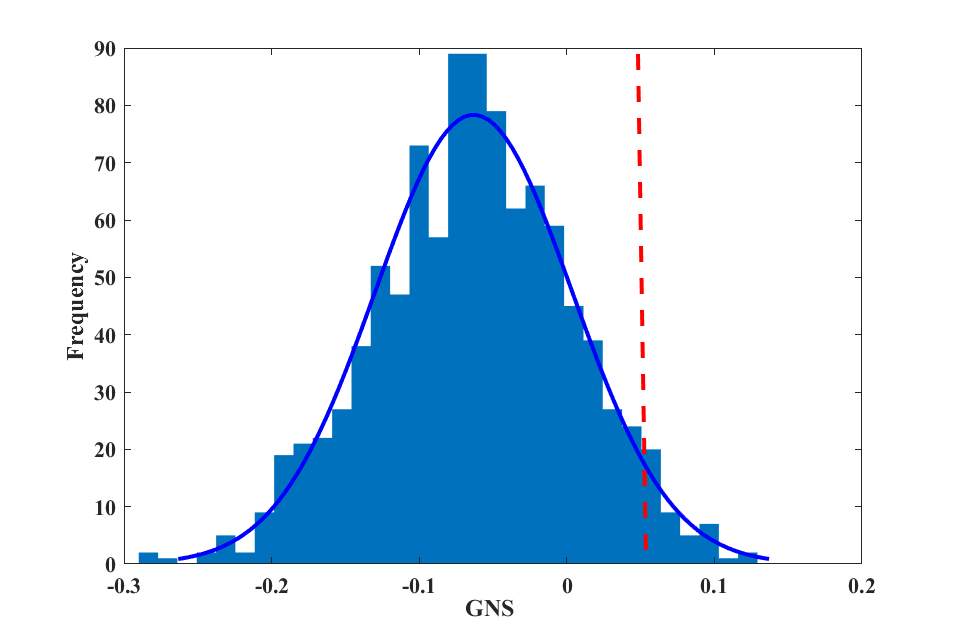

**Figure S2.** A permutation analysis confirmed that the enhanced intergroup coupling shown was group-specific. The figure showed the distribution of the permutated average intergroup coupling enhancement. The black lines indicated the positions of the true means of the original groups.
